# Feasibility of Using Functional Near-Infrared Spectroscopy for Characterizing Misophonia-Related Brain Activity

**DOI:** 10.64898/2026.08.30.748122

**Authors:** Shibo Zhou, Alaina Cunningham, Jane Miller-Laquerre, Benjamin Kirby, Yuanyuan Gao

## Abstract

**Significance:** Misophonia is characterized by decreased tolerance to specific sounds (“triggers”) that evoke negative emotional, physiological, and behavioral responses. To understand the neural mechanisms underlying misophonia, functional neuroimaging has been proposed such as functional magnetic resonance imaging (fMRI). However, the fMRI environment introduces substantial noise that increases anxiety, which is one of the comorbidities of misophonia, confounds auditory stimuli, and is costly. We therefore propose an alternative, functional near-infrared spectroscopy (fNIRS) to detect misophonia-related responses.

**Aim:** We evaluated the feasibility of fNIRS to detect misophonia-related sound-evoked responses.

**Approach:** We measured fNIRS, electrodermal activity (EDA), and photoplethysmography (PPG) data from ten adults with minimal to mild misophonia symptoms while they listened to trigger, unpleasant, and neutral sounds. Participants rated their annoyance and antisociality after each sound.

**Results:** Compared with neutral sounds, trigger and unpleasant sounds received higher subjective ratings and evoked greater neural response in the orofacial area. Neural responses tended to increase with annoyance ratings. Significantly lower heart rate (HR) was observed during unpleasant sounds than during neutral and trigger sounds, with no significant difference in EDA features.

**Conclusions:** fNIRS can measure misophonia responses in a quiet setting. Both brain and physiological responses have the same trend observed in prior fMRI studies, verifying its feasibility.

## 1 Introduction

Misophonia is characterized by symptoms of decreased tolerance to specific sounds or sound-related stimuli (“triggers”) that evoke strong negative emotional, physiological, and behavioral responses not observed in most people^1^. Common symptoms of misophonia include irritation, anger, disgust, muscular tension, and avoidance of situations in which triggers occur^1–3^. Moderate to severe symptoms of misophonia have been reported to affect a large portion of the population (6–20%)^4–6^. These symptoms affect the occupational, social, and domestic life of sufferers, who may change or quit jobs, become socially isolated, and experience strained family relationships^2,3,7,8^. Investigating neural mechanisms underlying misophonia symptoms is one way to help us understand misophonia. To achieve this, previous functional magnetic resonance imaging (fMRI) studies have reported abnormal activation of the orofacial motor cortex and functional connectivity across salience, auditory, and limbic networks^9–12^ related to misophonia.

However, the magnetic resonance imaging (MRI) environment is not ideal for studying auditory responses in misophonia due to substantial background noise (90-120 dB)^13,14^ that potentially contaminates auditory stimuli, making neutral sounds non-neutral and altering the perceptual qualities of both trigger and control stimuli. The enclosed MRI environment could also potentially heighten anxiety^15–19^ and hyperacusis responses, which are common comorbidities^20,21^ of misophonia.

Additionally, fMRI is not optimal for routine use because of its high cost, limited accessibility, and the need for participants to go inside an MRI scanner. For example, routine intervention facilitated by neuroimaging, such as neurofeedback, would be challenging to do using fMRI. However, neurofeedback could be effective in the treatment of misophonia since it has been shown to improve emotional regulation^22^, promote self-regulation of neural activity^23^, and reduce symptom severity across a range of neurological and psychiatric conditions^24–27^. Unlike pharmacological treatments, neurofeedback directly targets neural biomarkers associated with the disorder and enables participants to learn voluntary control over their own brain activity through repeated training. Because it is non-invasive and can be delivered over multiple training sessions, neurofeedback has the potential to produce durable therapeutic benefits with minimal side effects^28,29^. However, practical limitations make MRI unsuitable for routine clinical neurofeedback.

Functional near-infrared spectroscopy (fNIRS) is an alternative optical neuroimaging technique. fNIRS measures changes in oxygenated (HbO) and deoxygenated hemoglobin (HbR) concentrations in superficial cortical regions, providing an indirect assessment of local brain activation through neurovascular coupling^30–33^. The technique can be used during naturalistic tasks^34^ and combined with physiological recordings for multimodal measurements^35^. fNIRS has been applied to study neurodevelopment^36,37^, perception and cognition^38,39^, and psychiatric conditions including anxiety and depression^40,41^, which commonly co-occur with misophonia^21,42^. In contrast to fMRI, fNIRS is silent, portable, considerably less expensive, tolerant of natural movement, and deployable in outpatient clinics or even home-based settings, thus, being more suitable for misophonia research. To test the ability of fNIRS to detect misophonia-specific sound-related biomarkers, we replicated the experimental procedures used in prior fMRI work^12^, by using fNIRS instead of fMRI. We also simultaneously measured physiological responses, including photoplethysmography (PPG) and electrodermal activity (EDA). We determined whether the same neural activation, physiological responses, and subjective ratings were observed.

## 2 Methods

### 2.1 Participants

Twelve college students with minimal to mild misophonia symptoms were recruited for this study. Inclusion criteria were (1) age ≥18 years, (2) right-handedness, (3) no history of neurological disorders, (4) Misophonia Questionnaire (MQ)^4^ severity score < 7, and (5) normal hearing. We excluded left-handed people to keep consistency within the group regarding neuroimaging on both hemispheres. The MQ Severity Scale is a single item rated from 1 (minimal) to 15 (very severe); scores below 7 indicate minimal (1–3) or mild (4–6) misophonia symptoms, without clinically significant sound sensitivity or misophonia^4,43^. Normal hearing was confirmed by a bilateral screen (20 dB HL at octave intervals from 250 Hz to 8000 Hz). Two participants were excluded because of hearing loss, leaving 10 participants for further analysis. Demographic and misophonia symptom information is summarized in **Table 1**. The study protocol was approved by the Institutional Review Board of Wichita State University, and all participants provided written informed consent before data collection.

**Table 1.** Demographics and questionnaire scores.

|  |  |
| --- | --- |
| **Number of subjects (*N*)** | 10 |
| **Sex (female)** | 9 |
| **Age (mean ± SD)** | 22.7 ± 5.5 |
| **Race/ethnicity (*N*)** |  |
| White | 5 |
| Asian | 3 |
| American Indian/Alaskan Native | 2 |
| Other | — |
| **MQ<sup>4</sup> (symptoms + behavioral) (mean ± SD)** | 14.1 ± 7.0 |
| **MQ<sup>4</sup> (severity) (mean ± SD)** | 2.6 ± 1.6 |

### 2.2 Experimental Paradigm

To validate the ability of fNIRS to detect misophonia related brain responses to trigger sounds, we adapted the experimental paradigm from an fMRI study^12^ (see original paper for paradigm details). Here we briefly describe the study design (**Fig. 1**): three categories of sounds were displayed to the participants using a block design (**Table 2**): (1) trigger sounds, which evoke a misophonic reaction in individuals with misophonia (e.g., chewing); (2) unpleasant sounds, which are generally aversive but do not evoke a misophonic response (e.g., screaming); and (3) neutral sounds, which evoke neither misophonic responses nor general aversion (e.g., rain). Each sound category consisted of 14 distinct sounds, totaling 42 sound clips (details of the clips are in section ‘Audio Stimuli’). Each sound clip lasted 15 s, after which participants used continuous sliders, on a scale from 0 to 4, to give two ratings: “How annoying the sound was” and “How antisocial the sound was”. Ratings were collected during a randomized 8- to 12-s interstimulus interval (ISI). A progress bar indicated the remaining response time. If no responses were made before the ISI ended, the next sound clip displayed automatically. Participants completed five runs lasting ∼11 min each, comprising a total of 126 fully randomized trials (42 clips × 3 repetitions). Between runs, participants were given a self-paced break to minimize fatigue. Data was collected in a double-walled sound booth. fNIRS, EDA, and PPG signals were acquired continuously throughout each run.

**Fig. 1.**
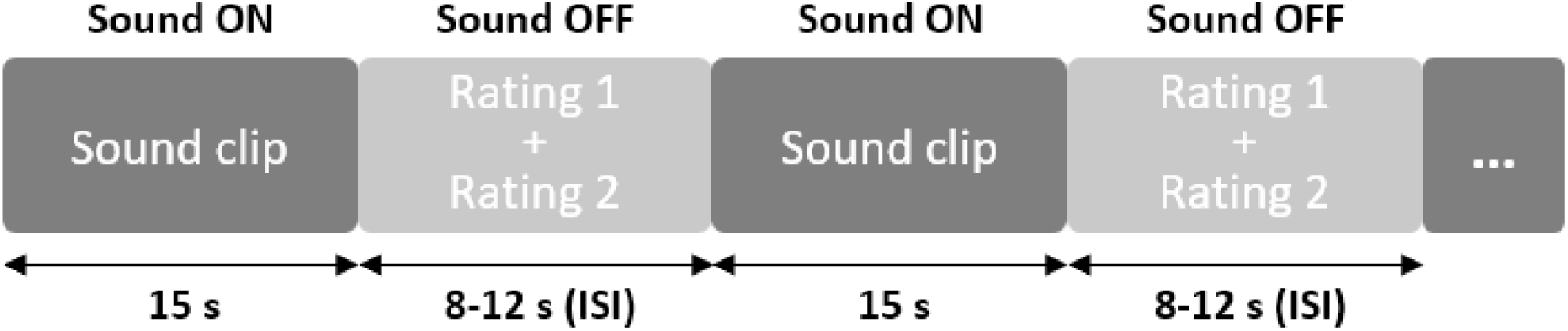
Experimental Design. Sound clips were displayed for 15 s (Sound ON). After each sound, participants used continuous sliders, on a scale from 0 to 4, to give two ratings: “How annoying the sound was” and “How antisocial the sound was” during a randomized 8- to 12-s interstimulus interval (ISI) (Sound OFF).

**Table 2.** List of sounds used in the experiment. ^9,12^ **a.** Recorded in a double-walled sound booth using a condenser microphone (Audio Technica AT2020) and digital audio interface (RME Fireface UCX). **b.** Selected and downloaded from SoundSnap^44^ online repository. *Trigger stimuli was previously used and showed efficacy in evoking misophonia symptoms in our previously published paper^45^.

| Trigger sounds<sup>a</sup> | Unpleasant sounds<sup>b</sup> | Neutral sounds<sup>b</sup> |
| --- | --- | --- |
| Apple crunching | Alarm sound | Brushing teeth |
| Breathing sound | Baby crying | Busy cafe |
| Chewing | Belch sound | Fan sound |
| Coughing and sniffing | Buzzer sound | Faucet sound |
| Crisps eating | Buzzing bees | Hair dryer sound |
| Cutlery sounds | Dentist drill | Helicopter sound |
| Eating food sound 1* | Female crying | Kettle boiling |
| Eating food sound 2* | Female scream | Phone ringing |
| Eating salad and cutlery | Jack hammer | Rain sound |
| Eating with slurping* | Male crying | Shower sound |
| Gulping water | Multiple dogs barking | Toilet flush |
| Packet opening and eating | Multiple infants crying | Traffic sound |
| Slurping* | Toddler crying | Vacuum cleaner |
| Sniffing* | Vomit sound | Washing machine |

### 2.3 Audio Stimuli

The audio stimuli were selected to match those used in Kumar et al.’s paper^12^ (Table 2). Since their audio stimuli were not open-access, we selected our neutral and unpleasant stimuli from an open-license online repository (SoundSnap^44^), and recorded misophonia trigger stimuli in a double-wall sound booth using a condenser microphone (Audio Technica AT2020) and digital audio interface (RME Fireface UCX). A subset of the trigger stimuli was used previously in a psychoacoustic study of misophonia^45^ and was selected to represent common categories of trigger sound that reliably evoke misophonia symptoms (Table 2). All stimuli were trimmed to 15 s overall length with 1 s linear ramps at the onset and offset. For the experiment, sound stimuli were presented at moderately-loud sound levels using insert earphones (EAR ER-3A).

### 2.4 fNIRS Measurements

We adapted an overlapping multi-distance fNIRS probe design from prior work^46^, with 7 sources and 16 detectors forming 50 measurement channels for each wavelength in each patch. We placed two patches over the left and right orofacial motor cortex regions resulting in a total of 100 channels for each wavelength. To differentiate HbO and HbR, we employed two wavelengths, 760 and 850 nm, resulting in a total of 200 measurement channels. The shorter source-detector separations were 19 mm apart, and the longer ones were 32.9 mm apart. We also placed two accelerometers at Cz and Pz locations to detect head movement. The configuration is illustrated in 2D probes (**Fig. 2a**), and 3D probes onto the cortical surface (**Fig. 2b**). The probe was designed in AtlasViewer^47^ and the cap was 3D printed using NinjaCap^48^. To ensure consistent optode placement across participants, we measured the Cz position of the head using the standard 10-20 international EEG locations and aligned it with the Cz marker of the cap. Two cascaded NIRSport2 16 × 16 devices (NIRx Medical Technologies LLC, Berlin, Germany) were used to record the fNIRS signal.

**Fig. 2.**
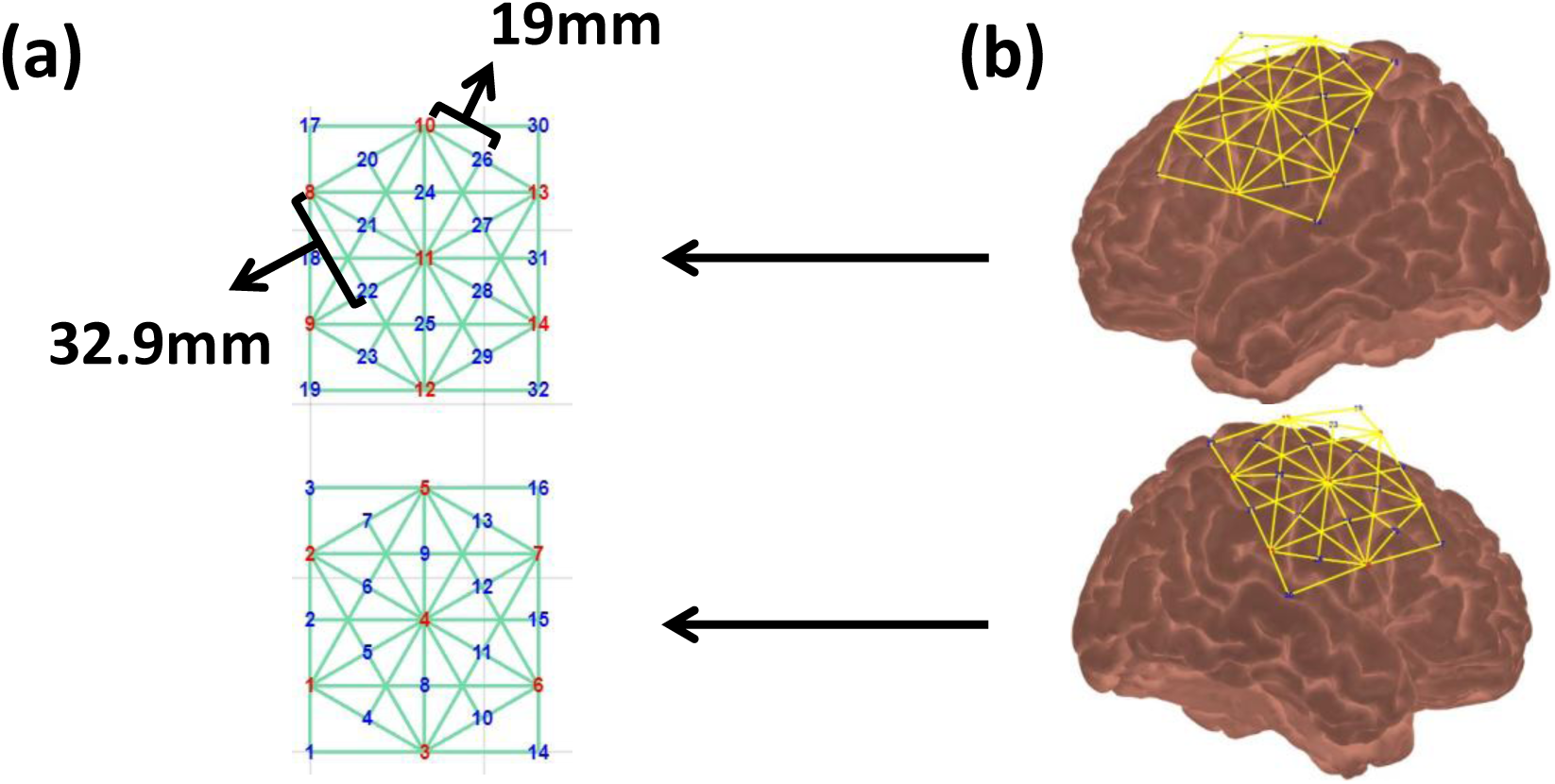
Probe Configuration. (a) 2D layouts probe of sources, detectors, and measurement channels. Blue numbers indicate detectors. Green lines indicate measurement channels. (b) 3D probe co-registration onto the brain atlas. Two patches over the left and right orofacial motor cortex regions. Yellow lines indicate the spatial arrangement of the channels relative to the underlying brain areas.

### 2.5 Physiological Measurements

To measure physiological changes during passive perception of sounds, PPG and EDA were continuously acquired during each run. These physiological signals were collected using a NIRxWINGS2 module integrated with the NIRSport2 system at 500 Hz (NIRx Medical Technologies, LLC, Berlin, Germany). The PPG sensor was clipped on the earlobe and the EDA sensors were attached to the middle and ring fingers of the left hand after applying conductive gel.

### 2.6 fNIRS Data Processing

fNIRS data were processed using customized MATLAB pipelines integrating functions from Homer 3^49^ and the NeuroDOT^50^ Toolbox for standardized preprocessing and task-specific analysis. First, channels with a signal-to-noise ratio value less than 5 were pruned from further analysis. Raw light intensity data were converted to optical density (OD). Motion artifacts were detected and corrected using spline interpolation combined with Savitzky-Golay filtering^51^, followed by a 0.5 Hz low-pass filter to remove high-frequency physiological noise (e.g., cardiac pulsations). Residual motion artifacts were detected per channel. Trials containing motion artifacts within the −2 to 20 s analysis window were rejected. OD was subsequently converted to concentrations of HbO and HbR using the modified Beer-Lambert law. Task-related hemodynamic response functions (HRF) were modeled using a general linear model (GLM) with a modified gamma basis function, solved by ordinary least squares. Short-separation regression was performed using the average of all short-separation channels, and polynomial drift correction was used to remove slow signal drift.

To generate sensitivity matrices, forward modeling of photon migration was performed via the MCXLAB^52^ package in AtlasViewer^47^. Image reconstruction was performed on both the brain and scalp, with Tikhonov regularization (α = 0.01)^53,54^, to generate estimates of peak HRF amplitude at each cortical vertex, for both HbO and HbR. All subsequent analyses only used HbO since HbR dynamics are highly correlated with those of HbO^55–57^ and have significantly lower amplitude than HbO^58^.

### 2.7 Physiological Data Processing

#### 2.7.1 PPG data processing

To assess stimulus-evoked physiological arousal from the PPG data, we extracted heart rate (HR) using the PhysioNet Cardiovascular Signal Toolbox^59^. The first 6 s of each recording were discarded to allow signals to stabilize. Next, we used an arterial blood pressure waveform algorithm^60^ to detect PPG wave peaks. We then visually inspected the PPG waveform with detected peaks to verify data quality. From PPG wave peaks, we calculated mean inter-beat interval (IBI) within each 15-s window. Finally, IBI was converted to HR (60/mean IBI, beats per minute).

#### 2.7.2 EDA data processing

To quantify stimulus-evoked physiological arousal from the EDA data, we extracted five distinct features: the number of significant skin conductance responses (nSCR), the sum of significant SCR amplitudes (AmpSum), the maximum phasic activity (PhasicMax), the time integral of the phasic driver (iSCR), and the tonic skin conductance (Tonic) following the procedure below.

We processed EDA data using the MATLAB Ledalab toolbox^61^, following a pipeline adapted from Loisel-Fleuriot et al^62^. The first 5 s of each recording were discarded to allow signals to stabilize, and the data were downsampled to 50 Hz to reduce computational load. Because physiologically relevant EDA signal is concentrated at low frequencies^63,64^, the signals were denoised with a 5-Hz low-pass Butterworth filter and then smoothed with an adaptive Gaussian kernel to remove residual noise. We decomposed the filtered signals into tonic (Skin Conductance Level; SCL) and phasic (Skin Conductance Response; SCR) components using Continuous Decomposition Analysis (CDA)^65^. Then we visually inspected SCR signals for each run to exclude runs lacking physiological responses (maximum stimulus-evoked amplitude < 0.05 µS in >50% of trials). Finally, features were extracted over a response window of 1-20 s after stimulus onset, with a minimum amplitude threshold of 0.01 µS, following recommendations for EDA analysis^63^.

### 2.8 Statistical Analysis

For the behavioral data, annoyance and antisocial ratings were averaged within each participant and sound category. To statistically evaluate differences across sound categories, we performed a one-way repeated-measures analysis of variance (RM-ANOVA) separately for the annoyance and antisocial ratings. The within-subject factor was sound category (trigger, unpleasant, neutral). Significant main effects were followed by Bonferroni-corrected pairwise comparisons to identify specific differences between sound categories (trigger vs. unpleasant, trigger vs. neutral, and unpleasant vs. neutral). Statistical significance was set at *p* < 0.05.

Separate Linear mixed-effects models^66,67^ were used to examine the effect of sound category (trigger, unpleasant, neutral) on brain activation (at each cortical vertex) and six physiological responses (nSCR, AmpSum, PhasicMax, iSCR, SCL and HR). Sound category was treated as a fixed effect. We included participant-level random intercepts to account for individual differences. The resulting t-values were plotted as vertex-wise t-statistic maps.

To visualize the relationship between brain activation and subjective ratings, we averaged HbO responses within each participant and sound category across vertices whose sensitivity exceeded the 0.01 threshold. Annoyance ratings were also averaged within each participant and sound category. Mean HbO responses were then plotted against the corresponding annoyance ratings, and an ordinary least-squares regression line was fitted to the data points.

## 3 Results

We collected 48 runs from the 10 participants. Two of the expected 50 runs were lost due to operator error.

### 3.1 Behavioral Results

We compared annoyance and antisocial ratings across sound categories. Prior to statistical analysis, the normality of the data distribution was assessed using the Lilliefors test^68^. Results indicated that ratings within all three sound categories were normally distributed (*p* > 0.05). Mauchly’s test indicated that the assumption of sphericity was not violated for both ANOVA models (annoyance and antisocial) (*p* > 0.05). The results are summarized in **Fig. 3**.

**Fig. 3.**
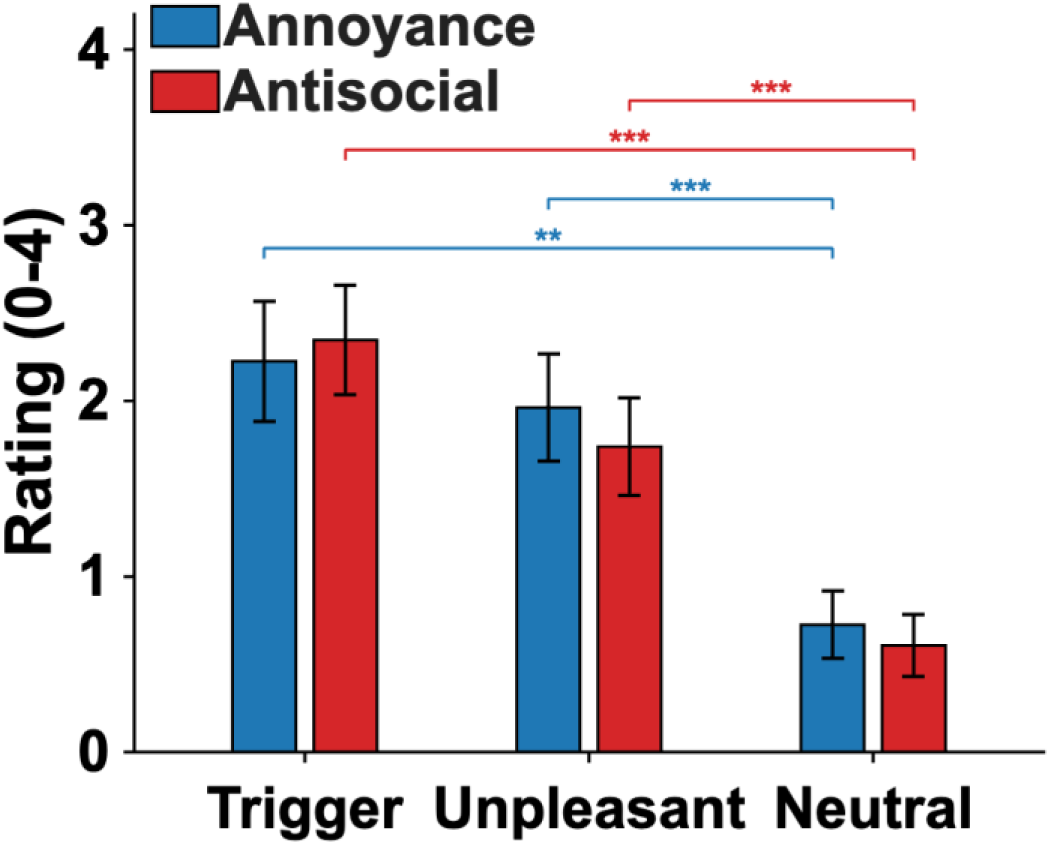
Comparisons of annoyance and antisocial ratings across three sound categories. Bar plots display annoyance (blue) and antisocial (red) ratings for trigger, unpleasant, and neutral sounds. Both annoyance and antisocial ratings were significantly lower for neutral sounds compared to trigger and unpleasant sounds, whereas no significant differences were observed between trigger and unpleasant sounds. Error bars represent ± standard error of the mean (SEM). Statistical significance between sound categories is indicated by brackets and asterisks (\**p* < 0.05, \*\**p* < 0.01, \*\*\**p* < 0.001; Bonferroni-corrected).

Sound category had a significant main effect on both annoyance ratings (*F*(2,18) = 34.36, *p* < 0.001) and antisocial ratings (*F*(2,18) = 26.27, *p* < 0.001). For both ratings, pairwise comparisons indicated that neutral sounds were rated significantly lower than trigger (annoyance: *p* = 0.001; antisocial: *p* < 0.001) and unpleasant sounds (annoyance: *p* < 0.001; antisocial: *p* < 0.001), with no significant difference between trigger and unpleasant sounds (annoyance: *p* = 0.701; antisocial: *p* = 0.077).

### 3.2 fNIRS Results

The group-averaged HbO and *t*-statistic maps, shown in **Fig. 4**, revealed activation over bilateral orofacial motor cortex for all three sound categories. Both HbO and t-statistic maps showed the largest activation for unpleasant sounds, particularly in the right hemisphere, followed by trigger sounds, with the smallest activation observed for neutral sounds.

**Fig. 4.**
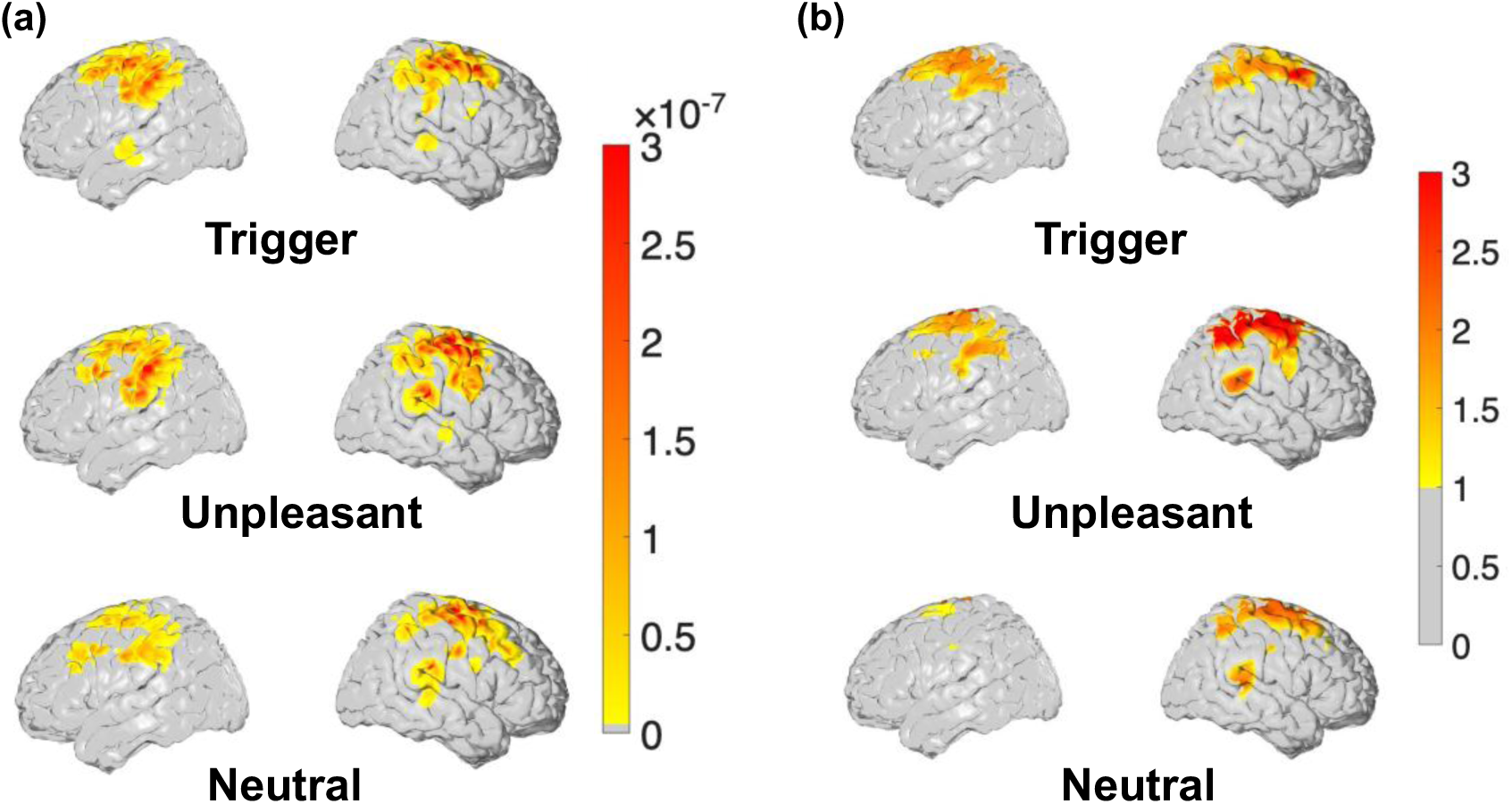
**Brain activation evoked by sound categories**. (a) Group-average HbO during each sound category. Color bar indicates HbO concentration (×10⁻⁷ M), with warmer colors (red) indicating higher HbO concentration. (b) T-statistic map during each sound category, derived from linear mixed-effects models. Color bar indicates t-statistic values, with warmer colors (red) indicating higher t-statistics. Gray vertices indicate t-statistics below 1.

**Fig. 5** shows the relationship between mean HbO responses and annoyance ratings. In the right orofacial motor cortex, HbO responses showed an increasing trend with annoyance ratings, although this trend was not statistically significant (*R*^2^ = 0.084, *p* = 0.120). Results for the left orofacial motor cortex are provided in Supplementary Material (**Fig. S1**).

**Fig. 5.**
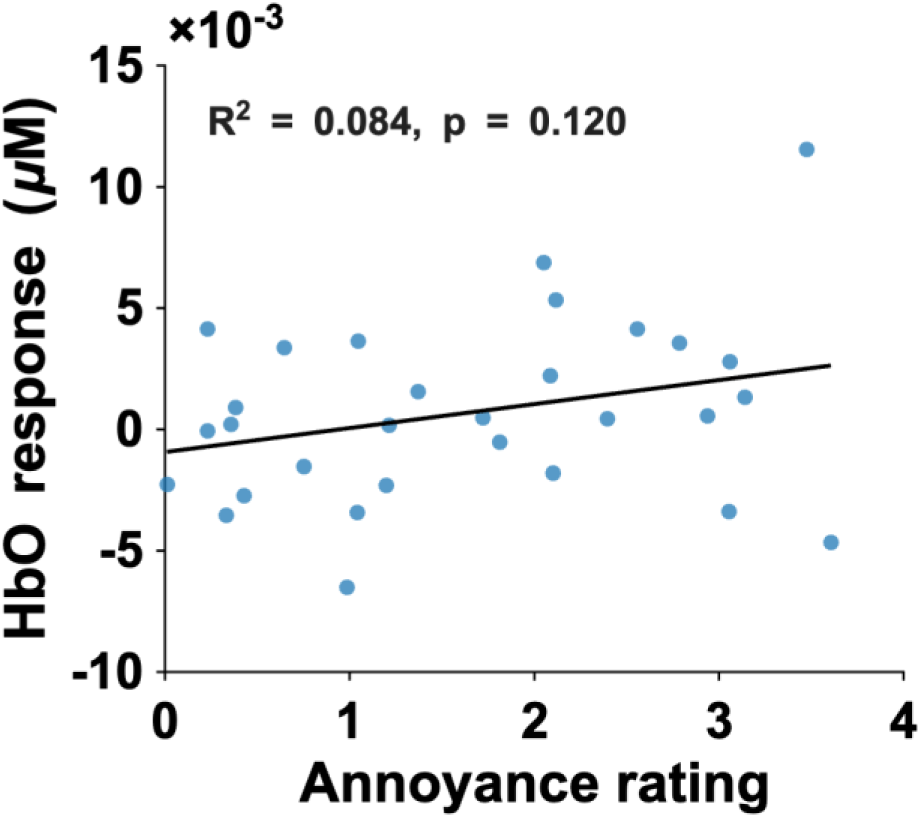
Scatter plot of mean HbO responses in the right orofacial motor cortex against annoyance ratings. HbO responses are expressed as concentration changes in μM. Each blue point represents one sound category in one participant. The solid line indicates the least-squares linear fit of the points (*R*² = 0.084, *p* = 0.120).

### 3.3 PPG and EDA results

For the PPG data analysis, all 48 runs passed visual inspection. HR was significantly lower during unpleasant sounds compared to both neutral sounds (*p* < 0.001) and trigger sounds (*p* < 0.01). No significant difference between trigger sounds and neutral sounds was observed (*p* = 0.22) (**Fig. 6**). HR time course responses to the three sound categories are shown in Supplementary Material **Fig. S11**.

**Fig. 6.**
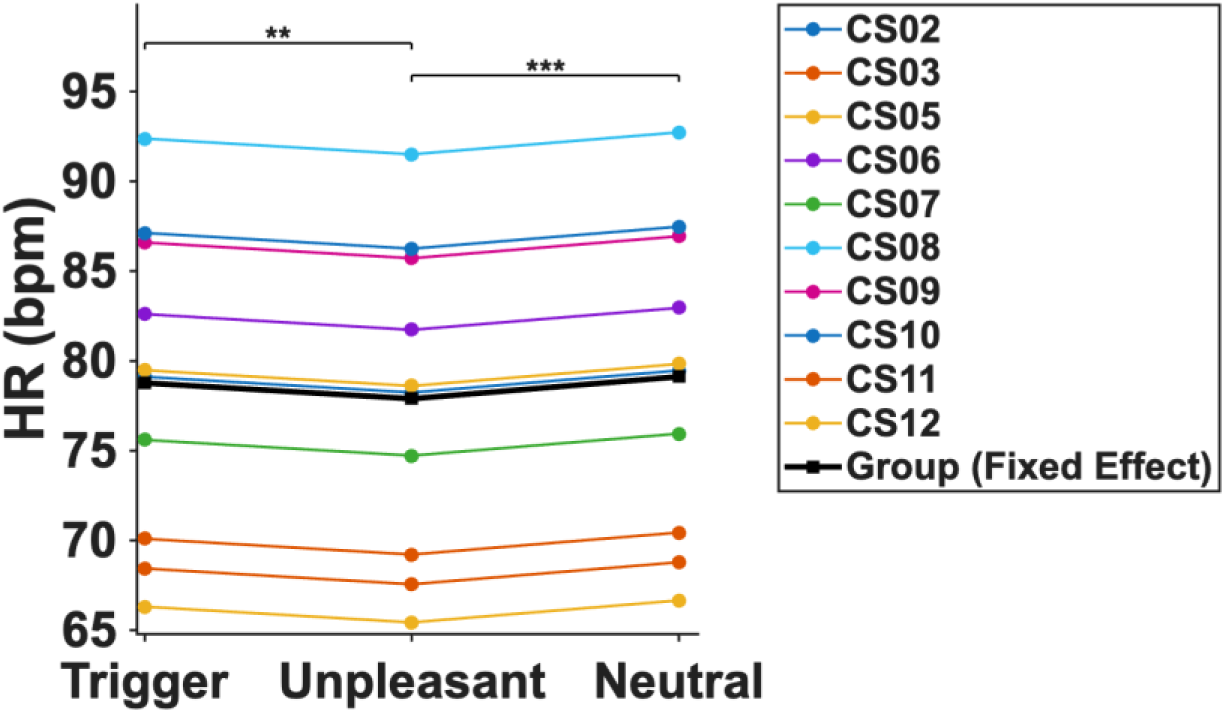
HR across sound categories. Thin colored lines represent individual HR across trigger, unpleasant and neutral sounds. Thick black line indicates the group-level fixed effect derived from the linear mixed-effects model. HR decreased significantly during unpleasant sounds compared to both neutral sounds (***p < 0.001) and trigger sounds (**p < 0.01). There was no significant change in HR between trigger sounds and neutral sounds.

For the EDA data analysis, we excluded 9 runs (distributed across three participants; Supplementary Material, **Fig. S2-S10**) due to poor signal quality, resulting in 39 runs for analysis. There was no significant effect of sound category on any of the five EDA features (all *p* > 0.05). EDA time course responses to the three sound categories are shown in Supplementary Material **Fig. S12**.

## 4 Discussion

We evaluated the ability of fNIRS to detect misophonia-specific sound-evoked hemodynamic responses in the orofacial motor cortex, using an experimental paradigm adapted from prior fMRI work^9,12^. Both trigger and unpleasant sounds were rated as more annoying and antisocial than neutral sounds. Brain activation followed the same pattern: trigger and unpleasant sounds evoked greater HbO responses than neutral sounds. In the right orofacial motor cortex, HbO responses showed an increasing trend with annoyance ratings, although this trend was not statistically significant. HR was significantly lower during unpleasant sounds than during trigger and neutral sounds, and none of the five EDA features differed across sound categories. The results were comparable to the prior fMRI work^9,12^, validating the feasibility of fNIRS.

Subjective ratings were consistent with the previous work^9^, with both trigger and unpleasant sounds being rated higher than neutral sounds. In our results, trigger sounds tended to be rated higher than unpleasant sounds, rather than lower in the previous work^9^. However, the trend was not significant in our study, and the previous work did not show statistical analysis in the control group. This difference in sound rating may reflect differences in testing environment: sound stimuli in our study were presented in a quiet sound booth, whereas those in the previous work^9^ were presented inside an MRI scanner with substantial background noise, which may have contaminated trigger sounds. Another reason could be that the age and sample size differed between the two studies: our participants were younger and fewer in number (mean 22.7 years, N = 10) than the control group in the previous work^9^ (mean 32.5 years, N = 22).

Using the same study protocol, we found that the brain activation patterns detected by fNIRS were consistent with the ones detected by fMRI^12^: orofacial motor cortex activation in their control group was greatest for unpleasant sounds, followed by trigger and then neutral sounds. This consistency supports the feasibility of using fNIRS to detect misophonia-specific sound-evoked activation in the orofacial motor cortex. The brain activation pattern may reflect engagement of the mirror neuron system, in which listening to an action-related sound evokes a “mirroring” motor representation of the action that produced the sound^69–72^. Since trigger sounds are commonly produced by orofacial movements such as eating and chewing, these sounds may therefore increase activity in the corresponding orofacial motor regions^12^. Many unpleasant sounds in our paradigm also involve orofacial actions, including crying, screaming, and vomiting, which may recruit similar motor representations.

The activation we detected extended beyond the orofacial motor region reported in the prior fMRI study^12^. We reviewed photographs and videos to confirm the caps were placed correctly. One possible explanation of this difference is that both trigger and unpleasant sounds are not limited to orofacial sounds. The unpleasant sounds include a broader range of aversive sources. Several trigger sounds, such as cutlery and package opening, involve hand actions rather than orofacial actions. These sounds may engage non-orofacial motor representations through “mirroring”, thus contributing to broader spatial activation^73^. This interpretation is consistent with prior evidence that “mirroring” motor representations may also involve finger-related motor and somatosensory regions^73,74^. Given the limited spatial resolution of fNIRS, we interpret this broader activation at the level of the sensorimotor cortex rather than assigning it to specific cortical subdivisions. Another possible explanation is the testing environment noted above: our fNIRS data were collected in a quiet sound booth, which may have allowed detection of broader neural activation that was present but undetected in the noise fMRI environment^75^.

HbO responses showed an increasing trend with annoyance ratings, although the trend was not statistically significant. This direction was consistent with that reported previously^12^. This consistency should be interpreted with caution. We used continuous rating scales, rather than the discrete rating scales used in the previous work^12^. In addition, the previous work plotted brain activation for each individual sound, whereas we averaged responses within each sound category for each participant.

For HR, our results differed from previous work^9^: whereas their HR trend showed no differences across sound categories, we found significantly lower HR during unpleasant sounds compared to both neutral and trigger sounds. The mean HR time courses showed a similar pattern (**Fig. S11**). However, this pattern was consistent with findings from the broader psychophysiology studies, in which unpleasant sounds elicited HR deceleration^76,77^. This deceleration may be attributed to the allocation of attentional resources during passive listening, with unpleasant sounds recruiting more attentional resources than other sounds and thereby producing larger drops in HR. This difference may also reflect the statistical analyses performed. The previous work^9^ tested the interaction between group and sound category using a 2 × 3 ANOVA, whereas we compared sound categories within only one group, using linear mixed-effects models with participant-level random intercepts to account for individual differences. In our data, the HR differences across sound categories were not detectable without these random intercepts showing that individual HR baseline is different in our sample. For EDA, our finding of no significant differences across sound categories was consistent with previous work^9^. However, the mean EDA time courses showed an increasing trend following the onset of unpleasant sounds compared to trigger and neutral sounds (**Fig. S12**), which may reflect greater autonomic arousal during unpleasant sound presentation^76^.

Even though the brain connectivity was also the main findings in previous fMRI studies^9–12,74^, the associated right planum temporale (PT)^11,12^, right secondary visual cortex (V2)^12^, and anterior insula areas^9^ were not surface areas and could not be reached by fNIRS. In addition, resting-state connectivity lacks specificity to trigger sounds and would also not be suitable for our future use of fNIRS for routine monitoring or neurofeedback. Thus, we only focused on the trigger sound elicited brain activation in this study.

There are two limitations to this study: the small number of participants and the inclusion of only those with minimal to mild misophonia symptoms. Based on the promising results derived from this feasibility study, we plan to conduct future studies to include individuals with clinically significant misophonia and age- and sex-matched controls to enable group comparisons. To have enough power to detect differences between the two groups, we plan to have more participants in both misophonia and control groups.

Together, these findings demonstrate the feasibility of using fNIRS to detect misophonia-specific sound-evoked hemodynamic responses in the orofacial motor cortex, with corresponding behavioral and physiological differences. fNIRS offers a practical way of studying misophonia-related responses in a quiet environment. These findings provide a methodological foundation for extending this paradigm to individuals with misophonia, enabling future group comparisons, and ultimately supporting the development of fNIRS-based interventions.

## Code and Data Availability

The custom MATLAB scripts developed for data processing and figure generation are available at https://github.com/Shibo3697/misophonia-behavioral-physio.git

## Acknowledgments

This work was supported by the Startup Fund and the Mutual Mentoring Award from Wichita State University.

## References

1. S. E. Swedo et al., “Consensus definition of misophonia: A Delphi study,” Front Neurosci 16, 841816 (2022) [doi:10.3389/fnins.2022.841816].

2. A. Schröder, N. Vulink, and D. Denys, “Misophonia: Diagnostic criteria for a new psychiatric disorder,” PLOS ONE 8(1), e54706, Public Library of Science (2013) [doi:10.1371/journal.pone.0054706].

3. R. Rouw and M. Erfanian, “A large-scale study of misophonia,” J Clin Psychol 74(3), 453–479 (2018) [doi:10.1002/jclp.22500].

4. M. S. Wu et al., “Misophonia: Incidence, phenomenology, and clinical correlates in an undergraduate student sample,” Journal of Clinical Psychology 70(10), 994–1007 (2014) [doi:10.1002/jclp.22098].

5. X. Zhou, M. S. Wu, and E. A. Storch, “Misophonia symptoms among Chinese university students: Incidence, associated impairment, and clinical correlates,” Journal of Obsessive-Compulsive and Related Disorders 14, 7–12 (2017) [doi:10.1016/j.jocrd.2017.05.001].

6. J. Naylor et al., “The prevalence and severity of misophonia in a UK undergraduate medical student population and validation of the amsterdam misophonia scale,” Psychiatr Q 92(2), 609–619 (2021) [doi:10.1007/s11126-020-09825-3].

7. I. Jager et al., “Misophonia: Phenomenology, comorbidity and demographics in a large sample,” PLoS One 15(4), e0231390 (2020) [doi:10.1371/journal.pone.0231390].

8. I. J. Jager et al., “Cognitive behavioral therapy for misophonia: A randomized clinical trial,” Depress Anxiety 38(7), 708–718 (2020) [doi:10.1002/da.23127].

9. S. Kumar et al., “The brain basis for misophonia,” Curr Biol 27(4), 527–533 (2017) [doi:10.1016/j.cub.2016.12.048].

10. A. Schröder et al., “Misophonia is associated with altered brain activity in the auditory cortex and salience network,” Sci Rep 9, 7542 (2019) [doi:10.1038/s41598-019-44084-8].

11. H. A. Hansen et al., “Selective disruption of salience-network anterior insula connectivity in misophonia: A disorder-specific neural signature,” Human Brain Mapping 47(3), e70468 (2026) [doi:10.1002/hbm.70468].

12. S. Kumar et al., “The motor basis for misophonia,” J. Neurosci. 41(26), 5762–5770, Society for Neuroscience (2021) [doi:10.1523/JNEUROSCI.0261-21.2021].

13. Z. H. Cho et al., “Analysis of acoustic noise in MRI,” Magn Reson Imaging 15(7), 815– 822 (1997) [doi:10.1016/s0730-725x(97)00090-8].

14. D. L. Price et al., “Investigation of acoustic noise on 15 MRI scanners from 0.2 T to 3 T,” J Magn Reson Imaging 13(2), 288–293 (2001) [doi:10.1002/1522-2586(200102)13:2<288::aid-jmri1041>3.0.co;2-p].

15. Z. M. Almutlaq, “Discussion of the causes, effect and potential methods of alleviating patient anxiety when undergoing magnetic resonance imaging (MRI),” The Egyptian Journal of Hospital Medicine 72(5), 4473–4477, Pan Arab League of Continuous Medical Education (2018) [doi:10.21608/ejhm.2018.9515].

16. D. Van Minde, L. Klaming, and H. Weda, “Pinpointing moments of high anxiety during an MRI examination,” Int.J. Behav. Med. (2013) [doi:10.1007/s12529-013-9339-5].

17. Z. Y. Hamd et al., “How different preparation techniques affect MRI-induced anxiety of MRI patients: A preliminary study,” Brain Sciences 13(3) (2023) [doi:10.3390/brainsci13030416].

18. I. Mutschler et al., “Who gets afraid in the MRI-scanner? Neurogenetics of state-anxiety changes during an fMRI experiment,” Neurosci Lett 583, 81–86 (2014) [doi:10.1016/j.neulet.2014.09.021].

19. M. R. Hanna et al., “Examining the role of emotion regulation, anger, and anxiety in misophonia: A network model,” PLoS One 20(8), e0329920 (2025) [doi:10.1371/journal.pone.0329920].

20. C. R. Brennan et al., “Misophonia and hearing comorbidities in a collegiate population,” Ear Hear 45(2), 390–399 (2024) [doi:10.1097/AUD.0000000000001435].

21. M. Z. Rosenthal et al., “Phenotyping misophonia: Psychiatric disorders and medical health correlates,” Front Psychol 13, 941898 (2022) [doi:10.3389/fpsyg.2022.941898].

22. J. Zaehringer et al., “Improved emotion regulation after neurofeedback: A single-arm trial in patients with borderline personality disorder,” Neuroimage Clin 24, 102032 (2019) [doi:10.1016/j.nicl.2019.102032].

23. V. Zotev et al., “Self-regulation of human brain activity using simultaneous real-time fMRI and EEG neurofeedback,” NeuroImage 85, 985–995 (2014) [doi:10.1016/j.neuroimage.2013.04.126].

24. Z. Zhao et al., “Real-time functional connectivity-informed neurofeedback of amygdala-frontal pathways reduces anxiety,” Psychother Psychosom 88(1), 5–15 (2019) [doi:10.1159/000496057].

25. K. D. Young et al., “Randomized clinical trial of real-time fMRI amygdala neurofeedback for major depressive disorder: Effects on symptoms and autobiographical memory recall,” Am J Psychiatry 174(8), 748–755 (2017) [doi:10.1176/appi.ajp.2017.16060637].

26. K. Patel et al., “Effects of neurofeedback in the management of chronic pain: A systematic review and meta-analysis of clinical trials,” Eur J Pain 24(8), 1440–1457, London, England (2020) [doi:10.1002/ejp.1612].

27. H. Gevensleben et al., “Is neurofeedback an efficacious treatment for ADHD? A randomised controlled clinical trial,” J Child Psychol Psychiatry 50(7), 780–789 (2009) [doi:10.1111/j.1469-7610.2008.02033.x].

28. J. Leem et al., “Effectiveness, cost-utility, and safety of neurofeedback self-regulating training in patients with post-traumatic stress disorder: A randomized controlled trial,” Healthcare (Basel) 9(10), 1351 (2021) [doi:10.3390/healthcare9101351].

29. V. Meisel et al., “Neurofeedback and standard pharmacological intervention in ADHD: A randomized controlled trial with six-month follow-up,” Biological Psychology 95, 116–125 (2014) [doi:10.1016/j.biopsycho.2013.09.009].

30. A. Villringer and U. Dirnagl, “Coupling of brain activity and cerebral blood flow: Basis of functional neuroimaging,” Cerebrovasc Brain Metab Rev 7(3), 240–276 (1995).

31. F. F. Jöbsis, “Noninvasive, infrared monitoring of cerebral and myocardial oxygen sufficiency and circulatory parameters,” Science 198(4323), 1264–1267, American Association for the Advancement of Science (1977) [doi:10.1126/science.929199].

32. D. A. Boas et al., “The accuracy of near infrared spectroscopy and imaging during focal changes in cerebral hemodynamics,” Neuroimage 13(1), 76–90 (2001) [doi:10.1006/nimg.2000.0674].

33. M. A. Franceschini et al., “Hemodynamic evoked response of the sensorimotor cortex measured noninvasively with near-infrared optical imaging,” Psychophysiology 40(4), 548– 560 (2003) [doi:10.1111/1469-8986.00057].

34. P. Pinti et al., “Using fiberless, wearable fNIRS to monitor brain activity in real-world cognitive tasks,” J Vis Exp(106), 53336 (2015) [doi:10.3791/53336].

35. A. von Lühmann et al., “Towards Neuroscience of the Everyday World (NEW) using functional near-infrared spectroscopy,” Curr Opin Biomed Eng 18, 100272 (2021) [doi:10.1016/j.cobme.2021.100272].

36. A. Cristia et al., “An online database of infant functional near infrared spectroscopy studies: a community-augmented systematic review,” PLoS One 8(3), e58906 (2013) [doi:10.1371/journal.pone.0058906].

37. Y. Gao et al., “Longitudinal changes in functional neural activation and sensitization during face processing in fragile X syndrome,” Biol Psychiatry 97(5), 499–506 (2025) [doi:10.1016/j.biopsych.2024.06.020].

38. P. Pinti et al., “The present and future use of functional near-infrared spectroscopy (fNIRS) for cognitive neuroscience,” Ann N Y Acad Sci 1464(1), 5–29 (2020) [doi:10.1111/nyas.13948].

39. R. J. Lawrence et al., “Cortical correlates of speech intelligibility measured using functional near-infrared spectroscopy (fNIRS),” Hear Res 370, 53–64 (2018) [doi:10.1016/j.heares.2018.09.005].

40. S. Hu et al., “Prefrontal cortex alterations in major depressive disorder, generalized anxiety disorder and their comorbidity during a verbal fluency task assessed by multi-channel near-infrared spectroscopy,” Psychiatry Res 306, 114229 (2021) [doi:10.1016/j.psychres.2021.114229].

41. A.-C. Ehlis et al., “Application of functional near-infrared spectroscopy in psychiatry,” Neuroimage 85 **Pt** **1**, 478–488 (2014) [doi:10.1016/j.neuroimage.2013.03.067].

42. M. Siepsiak et al., “Psychiatric and audiologic features of misophonia: Use of a clinical control group with auditory over-responsivity,” Journal of Psychosomatic Research 156, 110777 (2022) [doi:10.1016/j.jpsychores.2022.110777].

43. M. Z. Rosenthal et al., “Development and initial validation of the Duke misophonia questionnaire,” Front Psychol 12, 709928 (2021) [doi:10.3389/fpsyg.2021.709928].

44. “Professional Sound Effects - Royalty-Free SFX | Unlimited Downloads,” <https://www.soundsnap.com> (accessed 13 April 2026).

45. B. J. Kirby, A. Cunningham, and O. M. Zant, “Psychoacoustic assessment of misophonia,” JASA Express Lett. 5(9), 094401 (2025) [doi:10.1121/10.0039238].

46. Y. Gao et al., “Short-separation regression incorporated diffuse optical tomography image reconstruction modeling for high-density functional near-infrared spectroscopy,” Neurophoton. 10(02) (2023) [doi:10.1117/1.NPh.10.2.025007].

47. C. M. Aasted et al., “Anatomical guidance for functional near-infrared spectroscopy: AtlasViewer tutorial,” Neurophoton 2(2), 020801 (2015) [doi:10.1117/1.NPh.2.2.020801].

48. A. von Lühmann et al., “NinjaCap: A fully customizable and 3D printable headgear for functional near-infrared spectroscopy and electroencephalography brain imaging,” NPh 11(3), 036601, SPIE (2024) [doi:10.1117/1.NPh.11.3.036601].

49. T. J. Huppert et al., “Homer: A review of time-series analysis methods for near-infrared spectroscopy of the brain,” Appl. Opt. 48(10), D280 (2009) [doi:10.1364/AO.48.00D280].

50. A. T. Eggebrecht and J. P. Culver, “NeuroDOT: An extensible Matlab toolbox for streamlined optical functional mapping,” in Diffuse Optical Spectroscopy and Imaging VII (2019), paper 11074_26, p. 11074_26, Optica Publishing Group (2019) [doi:10.1117/12.2527164].

51. S. Jahani et al., “Motion artifact detection and correction in functional near-infrared spectroscopy: A new hybrid method based on spline interpolation method and Savitzky-Golay filtering,” Neurophotonics 5(1), 015003 (2018) [doi:10.1117/1.NPh.5.1.015003].

52. Q. Fang and S. Yan, “MCX Cloud—a modern, scalable, high-performance and in-browser Monte Carlo simulation platform with cloud computing,” JBO 27(8), 083008, SPIE (2022) [doi:10.1117/1.JBO.27.8.083008].

53. D. A. Boas and A. M. Dale, “Simulation study of magnetic resonance imaging-guided cortically constrained diffuse optical tomography of human brain function,” Appl Opt 44(10), 1957–1968 (2005) [doi:10.1364/ao.44.001957].

54. A. Custo et al., “Anatomical atlas-guided diffuse optical tomography of brain activation,” Neuroimage 49(1), 561–567 (2010) [doi:10.1016/j.neuroimage.2009.07.033].

55. T. Yamada, S. Umeyama, and K. Matsuda, “Separation of fNIRS signals into functional and systemic components based on differences in hemodynamic modalities,” PLOS ONE 7(11), e50271, Public Library of Science (2012) [doi:10.1371/journal.pone.0050271].

56. X. Cui, S. Bray, and A. L. Reiss, “Functional near infrared spectroscopy (NIRS) signal improvement based on negative correlation between oxygenated and deoxygenated hemoglobin dynamics,” Neuroimage 49(4), 3039–3046 (2010) [doi:10.1016/j.neuroimage.2009.11.050].

57. T. J. Huppert et al., “A temporal comparison of BOLD, ASL, and NIRS hemodynamic responses to motor stimuli in adult humans,” Neuroimage 29(2), 368–382 (2006) [doi:10.1016/j.neuroimage.2005.08.065].

58. G. Strangman et al., “Factors affecting the accuracy of near-infrared spectroscopy concentration calculations for focal changes in oxygenation parameters,” Neuroimage 18(4), 865–879 (2003) [doi:10.1016/s1053-8119(03)00021-1].

59. A. N. Vest et al., “An open source benchmarked toolbox for cardiovascular waveform and interval analysis,” Physiol Meas 39(10), 105004 (2018) [doi:10.1088/1361-6579/aae021].

60. J. X. Sun, “Cardiac output estimation using arterial blood pressure waveforms,” Thesis, Massachusetts Institute of Technology (2006).

61. M. Benedek and C. Kaernbach, “A continuous measure of phasic electrodermal activity,” Journal of Neuroscience Methods 190(1), 80–91 (2010) [doi:10.1016/j.jneumeth.2010.04.028].

62. L. Loisel-Fleuriot et al., “A pilot study investigating affective forecasting biases with a novel virtual reality-based paradigm,” Sci Rep 13(1), 9321, Nature Publishing Group (2023) [doi:10.1038/s41598-023-36346-3].

63. W. Boucsein, Electrodermal Activity, Springer Science & Business Media (2012).

64. G. Geršak, “Electrodermal activity - a beginner’s guide,” Electrotechnical Review 87, 175–182 (2020).

65. M. Benedek and C. Kaernbach, “Decomposition of skin conductance data by means of nonnegative deconvolution,” Psychophysiology 47(4), 647–658 (2010) [doi:10.1111/j.1469-8986.2009.00972.x].

66. V. J. Carey and Y.-G. Wang, “Mixed-effects models in S and S-Plus,” Journal of the American Statistical Association 96(455), 1135–1136 (2001) [doi:10.1198/jasa.2001.s411].

67. H. Swaminathan and J. Rogers, “Estimation Procedures For Hierarchical Linear Models,” in Multilevel Modeling of Educational Data, A. A. O’Connell and D. B. McCoach, Eds., p. 0, Emerald Publishing Limited (2008) [doi:10.1108/978-1-60752-729-920251018].

68. H. W. Lilliefors, “On the Kolmogorov-Smirnov test for normality with mean and variance unknown,” Journal of the American Statistical Association 62(318), 399–402 (1967) [doi:10.1080/01621459.1967.10482916].

69. G. Rizzolatti and L. Craighero, “The mirror-neuron system,” Annu Rev Neurosci 27, 169–192 (2004) [doi:10.1146/annurev.neuro.27.070203.144230].

70. M. Iacoboni and M. Dapretto, “The mirror neuron system and the consequences of its dysfunction,” Nat Rev Neurosci 7(12), 942–951 (2006) [doi:10.1038/nrn2024].

71. E. Kohler et al., “Hearing sounds, understanding actions: action representation in mirror neurons,” Science 297(5582), 846–848, New York, N.Y. (2002) [doi:10.1126/science.1070311].

72. G. Buccino et al., “Action observation activates premotor and parietal areas in a somatotopic manner: an fMRI study,” Eur J Neurosci 13(2), 400–404 (2001).

73. G. Caetano, V. Jousmäki, and R. Hari, “Actor’s and observer’s primary motor cortices stabilize similarly after seen or heard motor actions,” Proc Natl Acad Sci U S A 104(21), 9058–9062 (2007) [doi:10.1073/pnas.0702453104].

74. H. A. Hansen et al., “Neural evidence for non-orofacial triggers in mild misophonia,” Front Neurosci 16, 880759 (2022) [doi:10.3389/fnins.2022.880759].

75. N. Gaab, J. D. E. Gabrieli, and G. H. Glover, “Assessing the influence of scanner background noise on auditory processing. I. An fMRI study comparing three experimental designs with varying degrees of scanner noise,” Human Brain Mapping 28(8), 703–720 (2007) [doi:10.1002/hbm.20298].

76. A.-M. Brouwer et al., “Perceiving blocks of emotional pictures and sounds: Effects on physiological variables,” Front. Hum. Neurosci. 7, Frontiers (2013) [doi:10.3389/fnhum.2013.00295].

77. K. Hume and M. Ahtamad, “Physiological responses to and subjective estimates of soundscape elements,” Applied Acoustics 74(2), 275–281 (2013) [doi:10.1016/j.apacoust.2011.10.009].

