## Supplementary Material for "Feasibility of Using Functional Near-Infrared Spectroscopy for Characterizing Misophonia-Related Brain Activity"

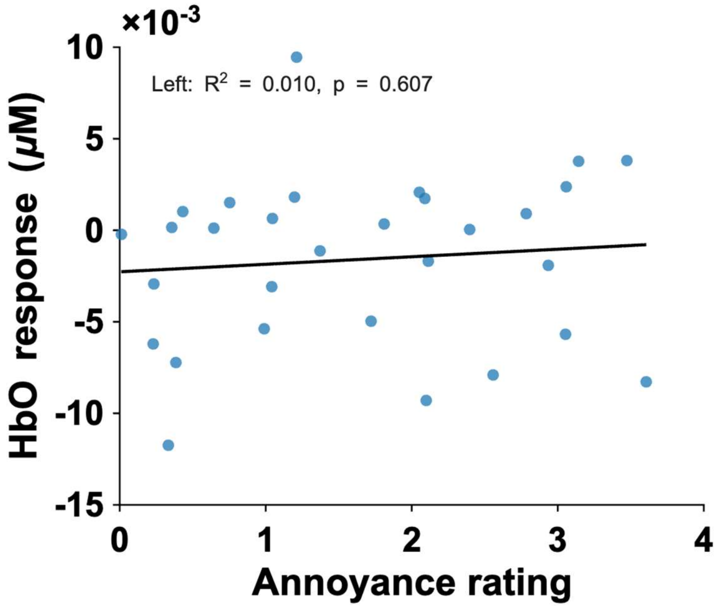


**Figure S1.** Scatter plot of mean HbO responses in the left orofacial motor cortex against annoyance ratings. HbO responses are reported in μM. Each blue point represents one sound category in one participant. The solid line indicates the least-squares linear fit of the points ($R^2 = 0.01$, $p = 0.607$).

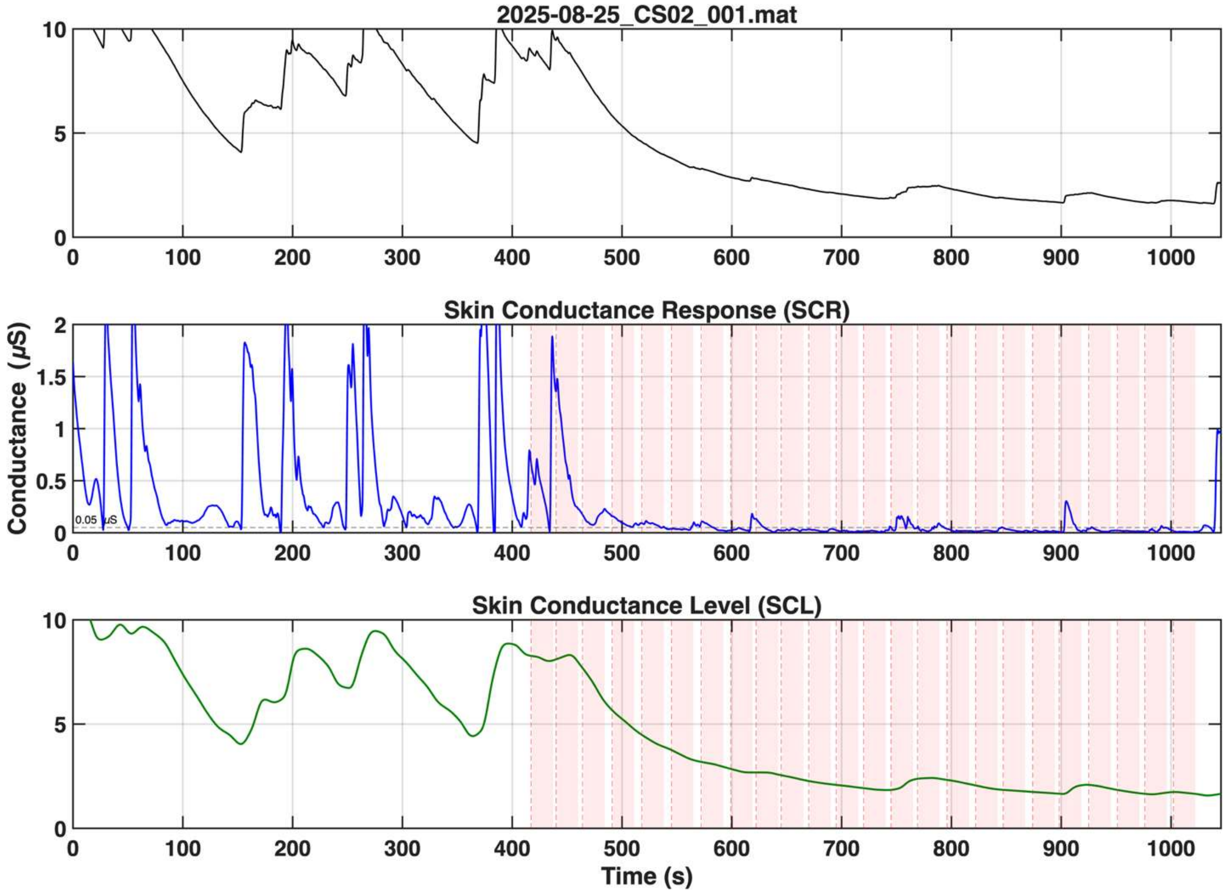

2025-08-25_CS02_001.mat
Skin Conductance Response (SCR)
Conductance (μS)
0.05 μS
Skin Conductance Level (SCL)
Time (s)

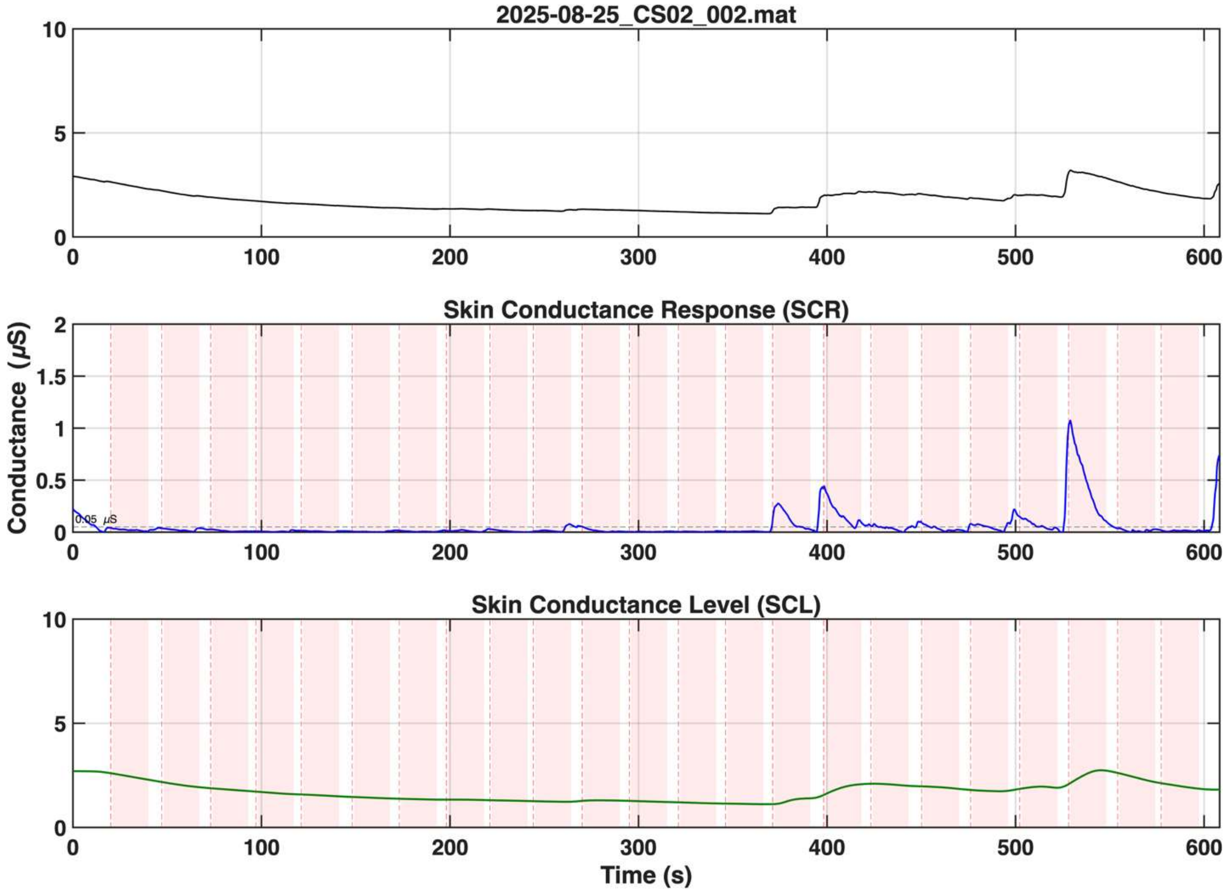
2025-08-25_CS02_002.mat
Skin Conductance Response (SCR)
Conductance (μS)
0.05 μS
Skin Conductance Level (SCL)
Time (s)

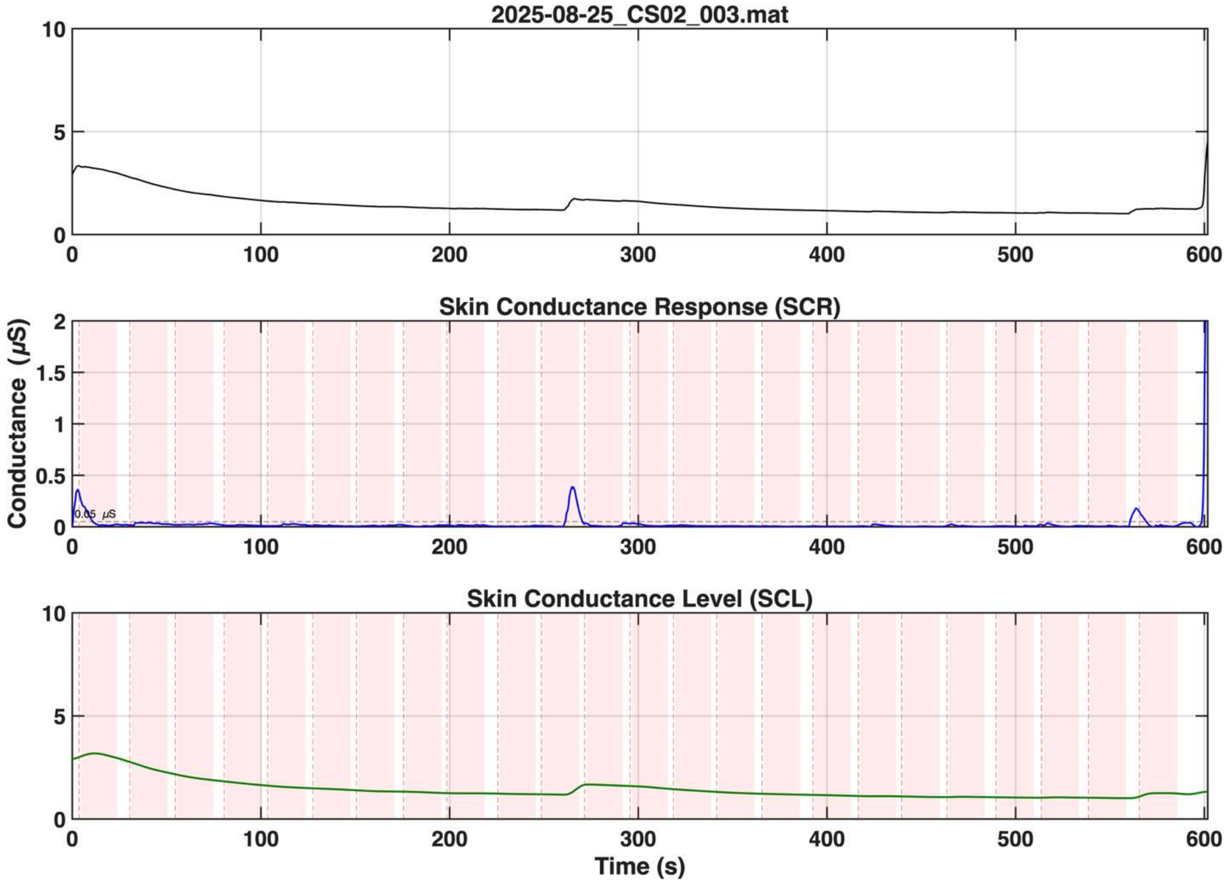

2025-08-25_CS02_003.mat
Skin Conductance Response (SCR)
Conductance (μS)
0.05 μS
Skin Conductance Level (SCL)
Time (s)

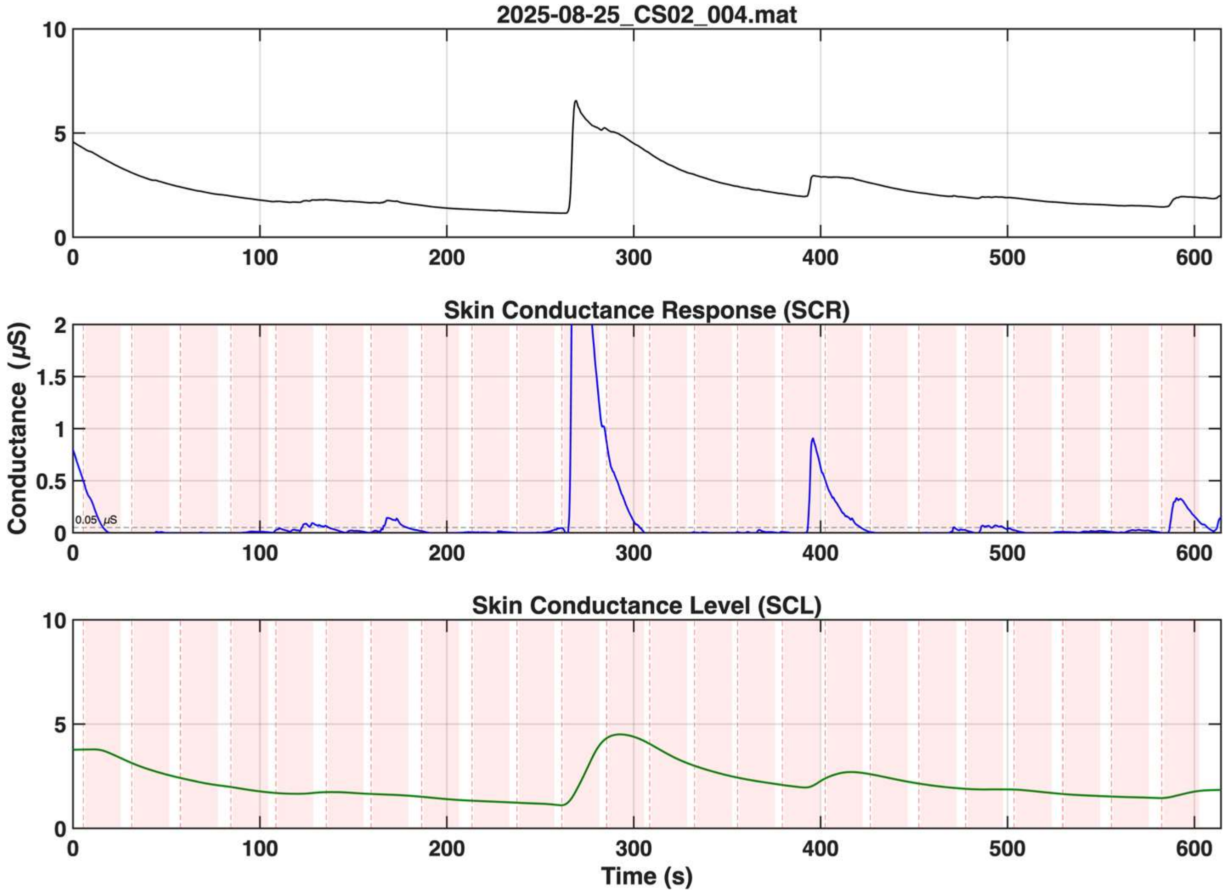

2025-08-25_CS02_004.mat
10
5
0
0
100
200
300
400
500
600
Skin Conductance Response (SCR)
Conductance (μS)
2
1.5
1
0.5
0
0.05 μS
Skin Conductance Level (SCL)
Time (s)

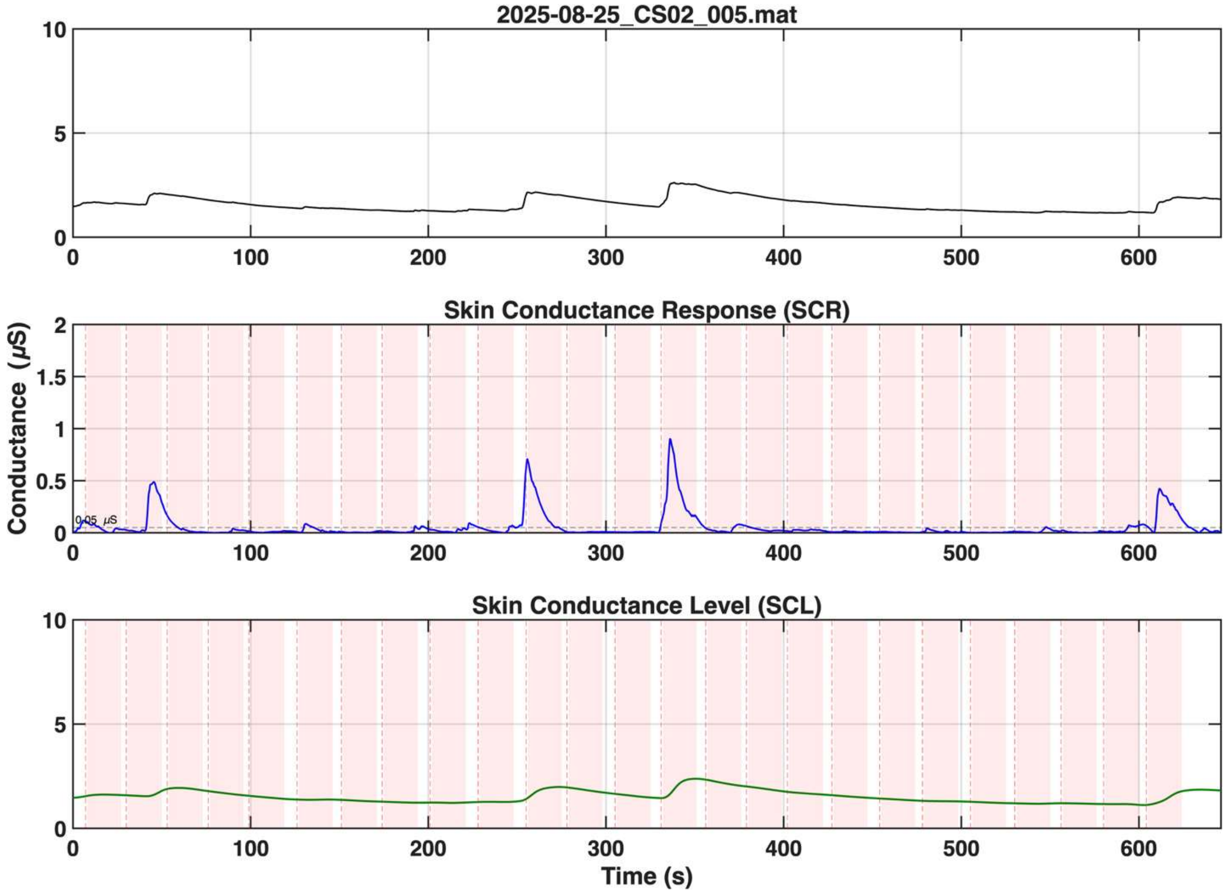

2025-08-25_CS02_005.mat
Skin Conductance Response (SCR)
Conductance (μS)
0.05 μS
Skin Conductance Level (SCL)
Time (s)

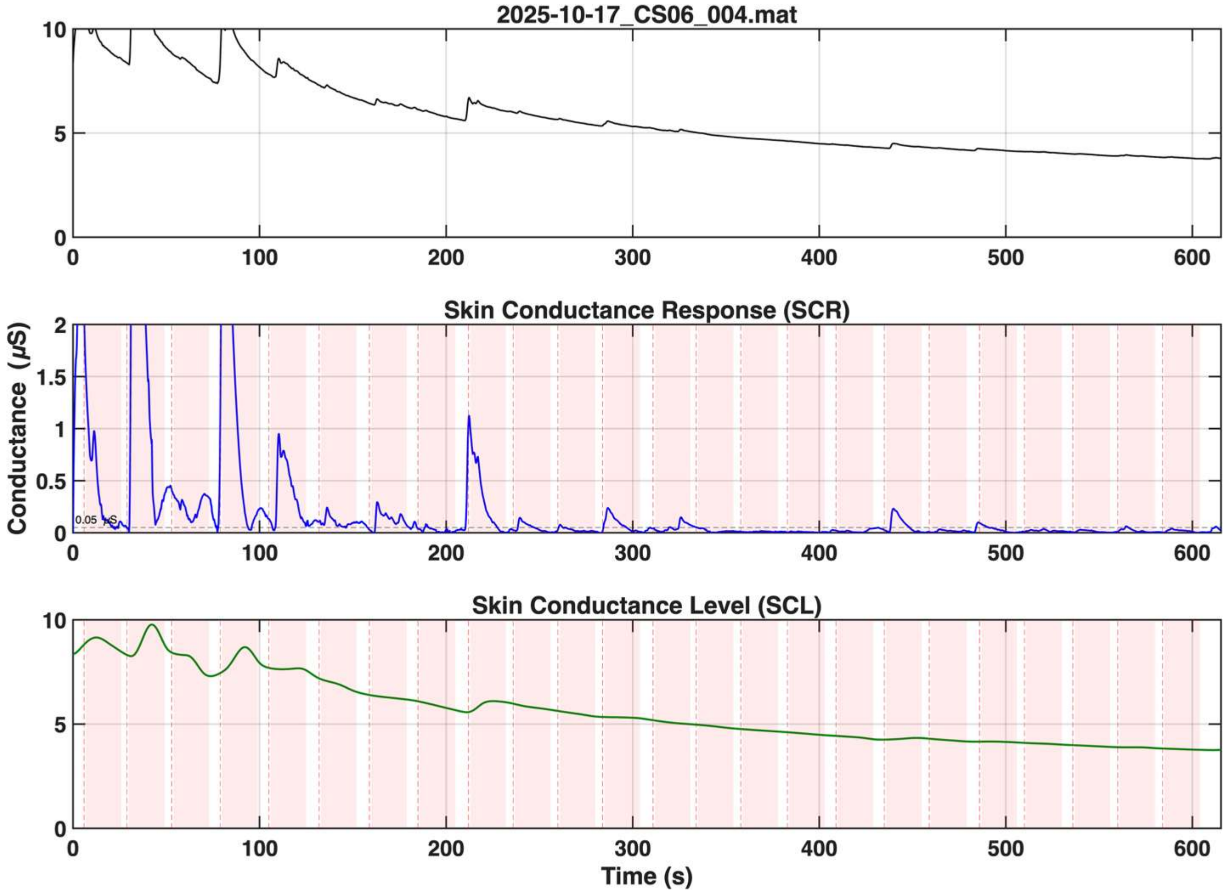

2025-10-17_CS06_004.mat
Skin Conductance Response (SCR)
Conductance (μS)
0.05 μS
Skin Conductance Level (SCL)
Time (s)

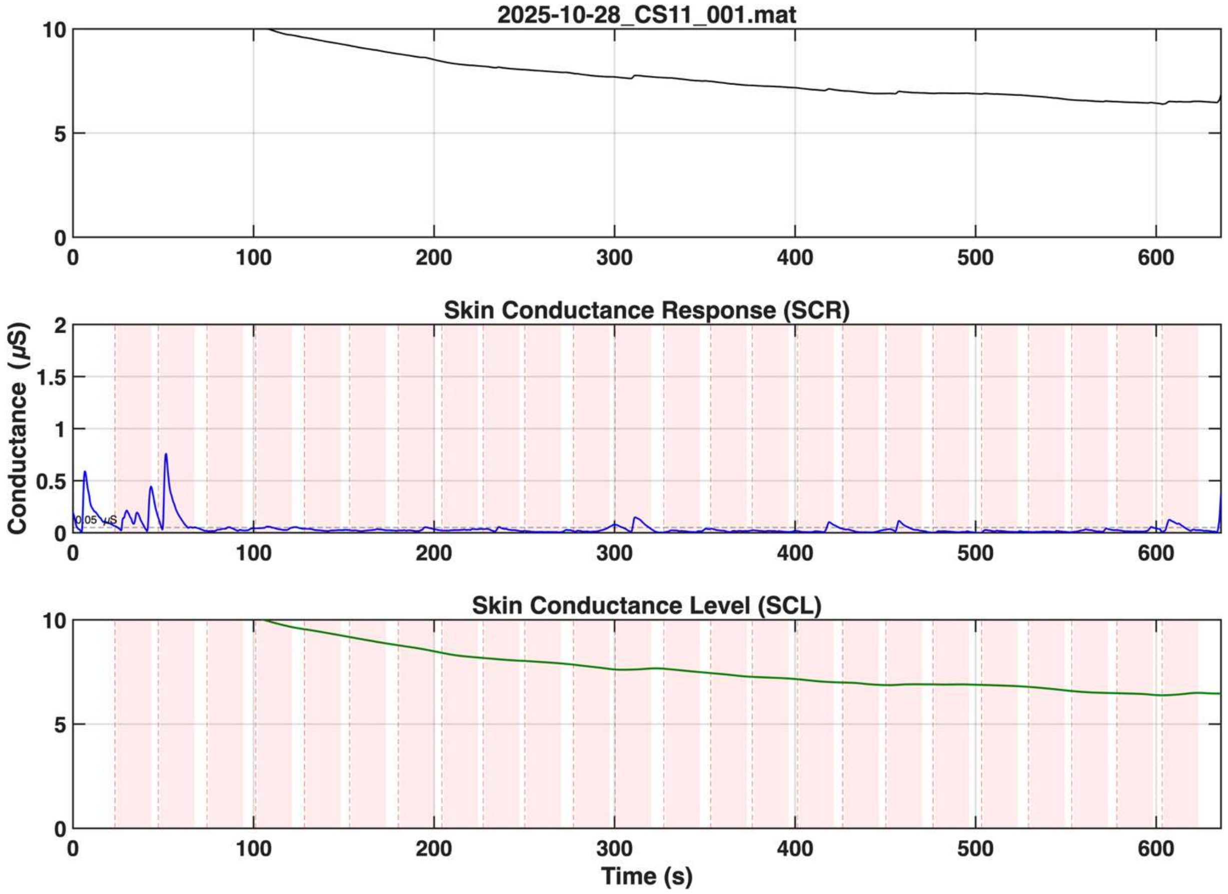

2025-10-28_CS11_001.mat
Skin Conductance Response (SCR)
Conductance (μS)
0.05 μS
Skin Conductance Level (SCL)
Time (s)

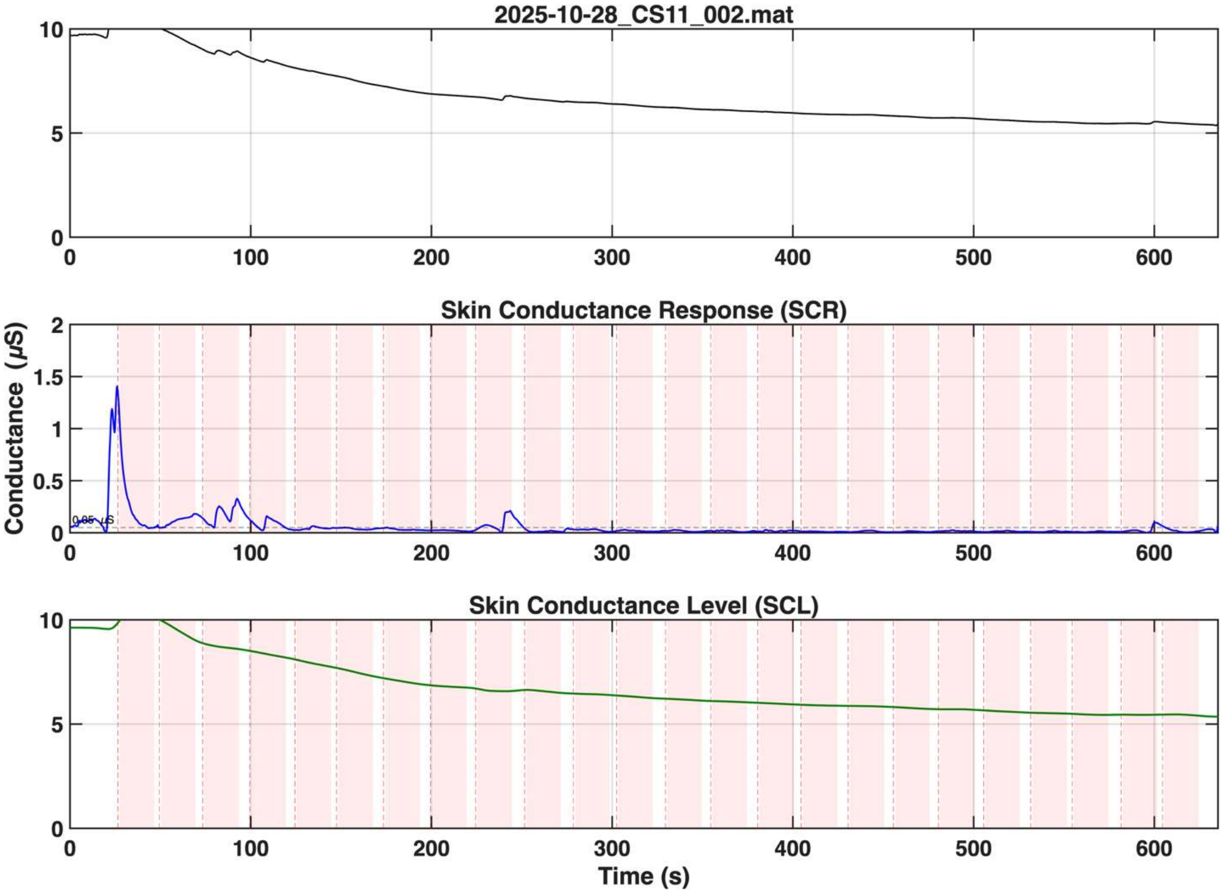

2025-10-28_CS11_002.mat
Skin Conductance Response (SCR)
Conductance (μS)
0.05 μS
Skin Conductance Level (SCL)
Time (s)

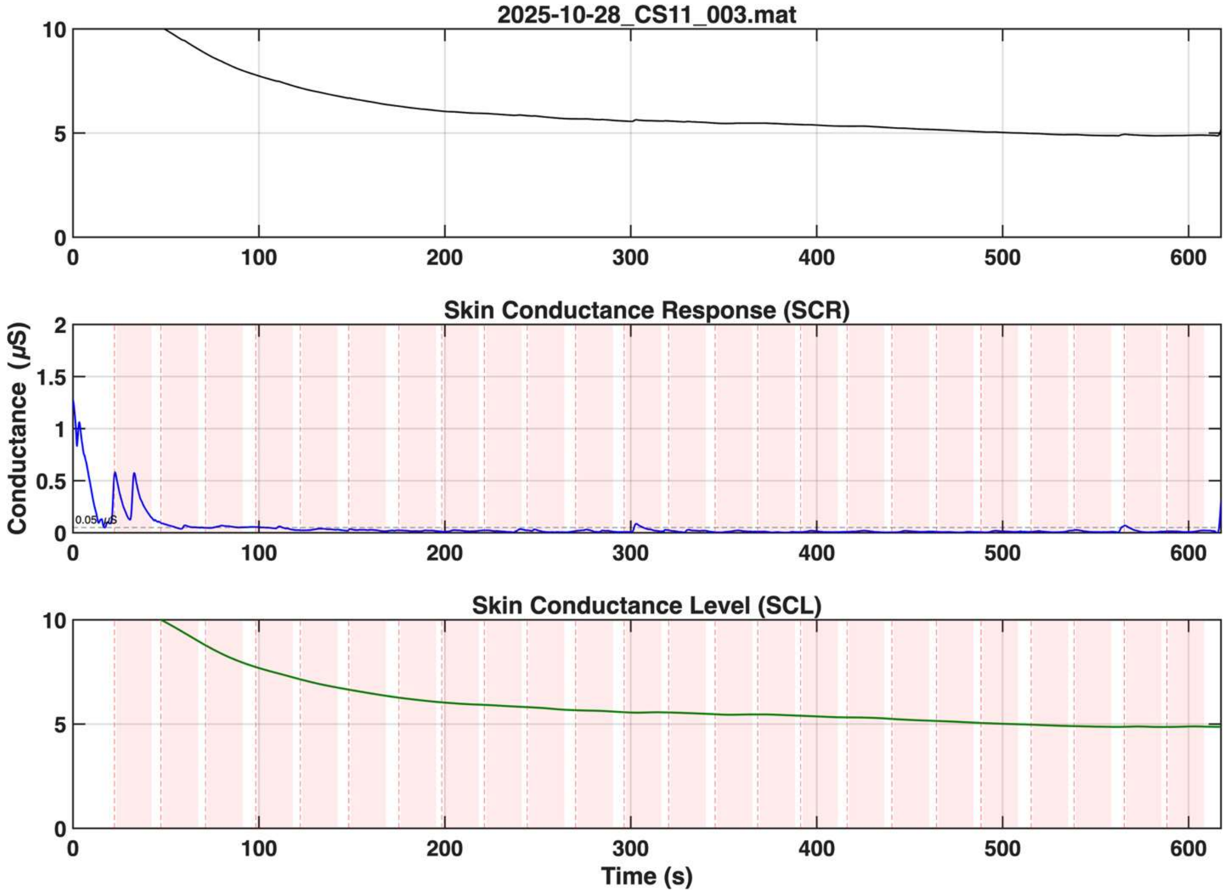


**Figure S2-S10. Electrodermal activity (EDA) runs excluded during visual inspection.** Each figure shows the raw conductance (top), the skin conductance response (SCR) (middle), and the skin conductance level (SCL) (bottom) extracted via Continuous Decomposition Analysis (CDA). These runs were excluded due to low physiological responses. Specifically, the maximum stimulus-evoked amplitude remained below 0.05 µS in over 50% of the trials within the SCR data in each of these runs.

**EDA and HR Visualization**

To allow comparison with previous work, the preprocessed EDA and heart rate (HR) signals were epoched for each trial from 5 s before sound onset to 20 s after onset, and baseline-corrected by subtracting the mean of the 5-s pre-stimulus period. For EDA, we used the preprocessed skin-conductance signal without further processing. For HR, we calculated instantaneous HR from each inter-beat interval (IBI). We excluded HR values below 45 bpm or above 120 bpm, and then spline-interpolated the remaining values onto a 10 Hz time course. The resulting EDA and HR epochs were averaged within each participant and sound category, and then averaged across participants.

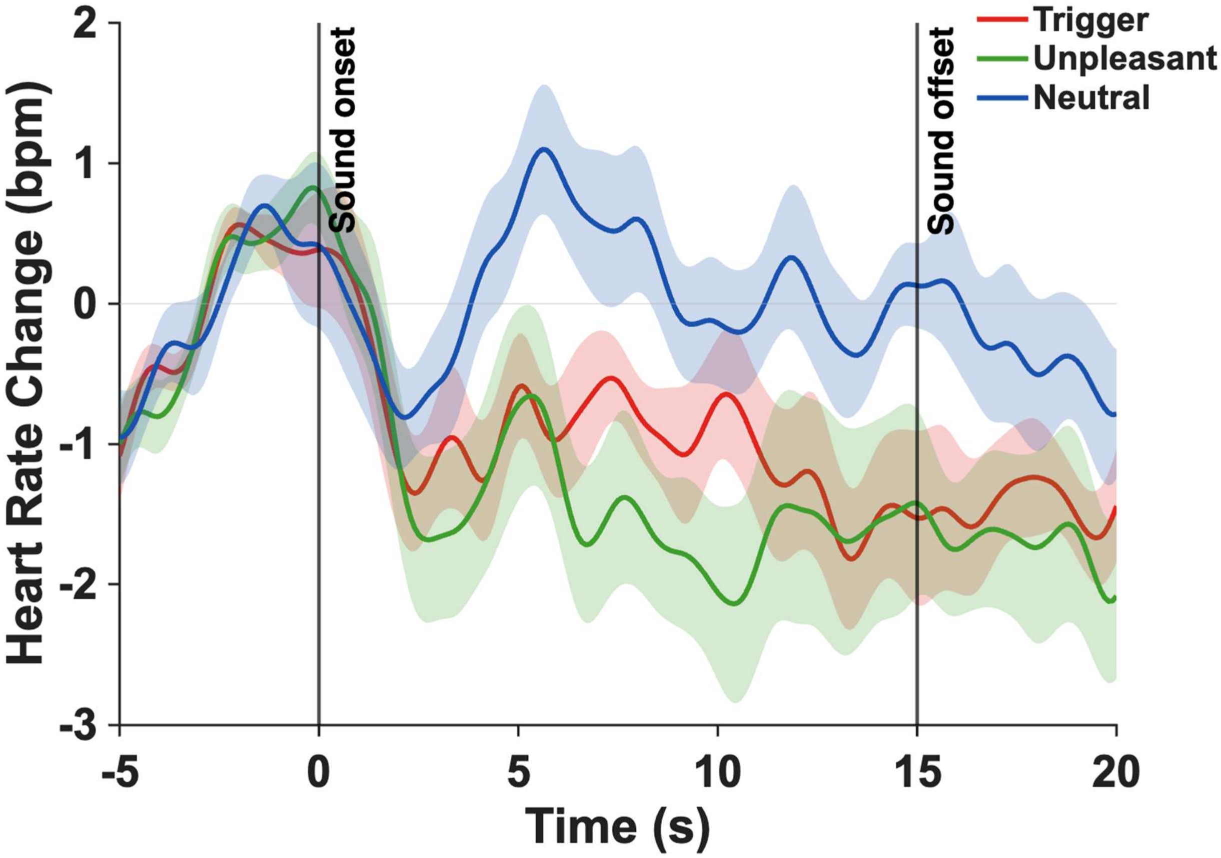


**Fig. S11 HR responses to the three sound categories.** Solid lines show the group mean. Shaded areas show the standard error of the mean across participants. Vertical lines mark sound onset (0 s) and offset (15 s).

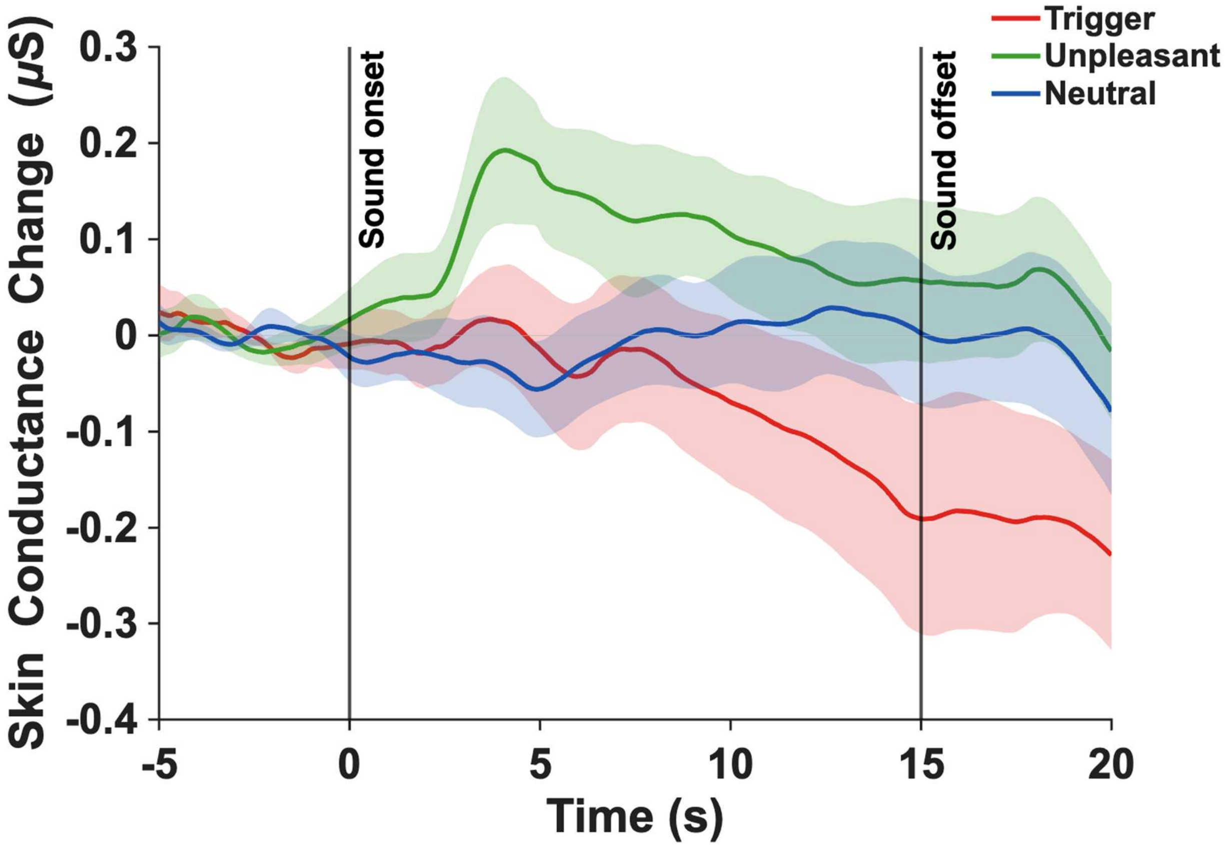


**Fig. S12 EDA responses to the three sound categories.** Solid lines show the group mean. Shaded areas show the standard error of the mean across participants. Vertical lines mark sound onset (0 s) and offset (15 s).
